# Meta-analysis of 1,200 transcriptomic profiles identifies a prognostic model for pancreatic ductal adenocarcinoma

**DOI:** 10.1101/355602

**Authors:** Vandana Sandhu, Knut Jorgen Labori, Ayelet Borgida, Ilinca Lungu, John Bartlett, Sara Hafezi-Bakhtiari, Rob Denroche, Gun Ho Jang, Danielle Pasternack, Faridah Mbaabali, Matthew Watson, Julie Wilson, Elin H. Kure, Steven Gallinger, Benjamin Haibe-Kains

**Author notes:** **Corresponding author:** Benjamin Haibe-Kains, 101 College Street, PMCRT 11-310, M5G1L7, Toronto, Canada.

## Abstract

**Background:** With a dismal 8% median 5-year overall survival (OS), pancreatic ductal adenocarcinoma (PDAC) is highly lethal. Only 10-20% of patients are eligible for surgery, and over 50% of these will die within a year of surgery. Identify molecular predictors of early death would enable the selection of PDAC patients at high risk.

**Methods:** We developed the Pancreatic Cancer Overall Survival Predictor (PCOSP), a prognostic model built from a unique set of 89 PDAC tumors where gene expression was profiled using both microarray and sequencing platforms. We used a meta-analysis framework based on the binary gene pair method to create gene expression barcodes robust to biases arising from heterogeneous profiling platforms and batch effects. Leveraging the largest compendium of PDAC transcriptomic datasets to date, we show that PCOSP is a robust single-sample predictor of early death (≤1 yr) after surgery in a subset of 823 samples with available transcriptomics and survival data.

**Results:** The PCOSP model was strongly and significantly prognostic with a meta-estimate of the area under the receiver operating curve (AUROC) of 0.70 (P=1.9e-18) and hazard ratio (HR) of 1.95(1.6-2.3) (P=2.6e-16) for binary and survival predictions, respectively. The prognostic value of PCOSP was independent of clinicopathological parameters and molecular subtypes. Over-representation analysis of the PCOSP 2619 gene-pairs (1070 unique genes) unveiled pathways associated with Hedgehog signalling, epithelial mesenchymal transition (EMT) and extracellular matrix (ECM) signalling.

**Conclusion:** PCOSP could improve treatment decision by identifying patients who will not benefit from standard surgery/chemotherapy and may benefit from alternate approaches.

**Abbreviations:**

AUROC: Area under the receiver operating curve
GO: Gene annotation
OS: Overall survival
PCOSP: Pancreatic cancer overall survival predictor
PDAC: Pancreatic ductal adenocarcinoma
TSP: Top scoring pairs.

## Introduction

Pancreatic ductal adenocarcinoma (PDAC) is a highly lethal malignancy with 5-year overall survival rate less than 8%^1^. The majority of patients (> 80%) are inoperable due to locally advanced or metastatic disease at time of diagnosis. While surgical resection is the key to curative treatment, it rarely results in long-term survival^2^. Hence, completion of multimodality treatment - surgery combined with adjuvant or neoadjuvant chemotherapy - is the standard of care for treatment of PDAC. However, even after surgical resection with curative intent, median survival does not exceed 28 months and half of those who undergo surgery develop recurrent disease, and die within a year after surgery^3,4^.Therefore, there is a need for a robust prognostic model to identify patients with high risk of early death based on molecular profiles of their tumors. Such a prognostic model would assist clinicians in identifying patients who might not benefit from surgery and standard adjuvant chemotherapy and may benefit from alternative treatment strategies.

Various clinical factors are prognostic following PDAC surgery such as lymph node metastasis status^5^, tumor grade^6^, margins^7^, degree of differentiation^8^ and protein biomarker *CA-19-9*^9^. However, the prognostic value of these clinical variables are insufficient to accurately stratify patients based on risk of disease recurrence^10,11^. With the advent of high-throughput next-generation molecular profiling technologies, multiple studies have released transcriptomic profiles of PDAC to the public domain. These gene expression profiles have been leveraged to identify molecular subtypes of PDACs^12–16^. While overlap between these subtypes^15^ supports the biological relevance of these published classification schemes^15^, they have not been designed to optimize prognostic value.

Previously published prognostic models were developed from small number of samples lacking proper validation in multiple datasets^17–21^. Attempts have been made recently to build a prognostic gene signature using pooled samples from multiple cohorts to identify patients at high risk of short-term survival post surgery^22–24^. However, they used samples profiled using either array or sequencing based method as the learning cohort, therefore the classifiers may perform better for subjects whose samples were profiled using only one of the two platforms.

To address these issues, we took advantage of a unique set of 89 PDAC profiled using both microarray and sequencing technologies to develop the Pancreatic Cancer Overall Survival Predictor (PCOSP) model. Using an independent set of PDAC transcriptomic profiles from 823 primary resected patients, we show that PCOSP is a robust single-sample predictor of early death (≤1 yr) after surgery with a meta-estimate of the area under the receiver operating characteristics curve of 0.70 (p=1.9e-18). We also show that PCOSP is significantly prognostic (meta-estimate of hazard ratio of 1.95; p=2.6e-16). Furthermore, we show that PCOSP performs significantly better than published prognostic models across microarray and sequencing datasets (Superiority test, P < 0.01). Our results support PCOSP as a potential tool to assist clinicians in decision making.

## Materials and Methods

The meta-analysis pipeline used to develop the PCOSP model and evaluate its prognostic value is provided in Figure 1.

**Figure 1.**
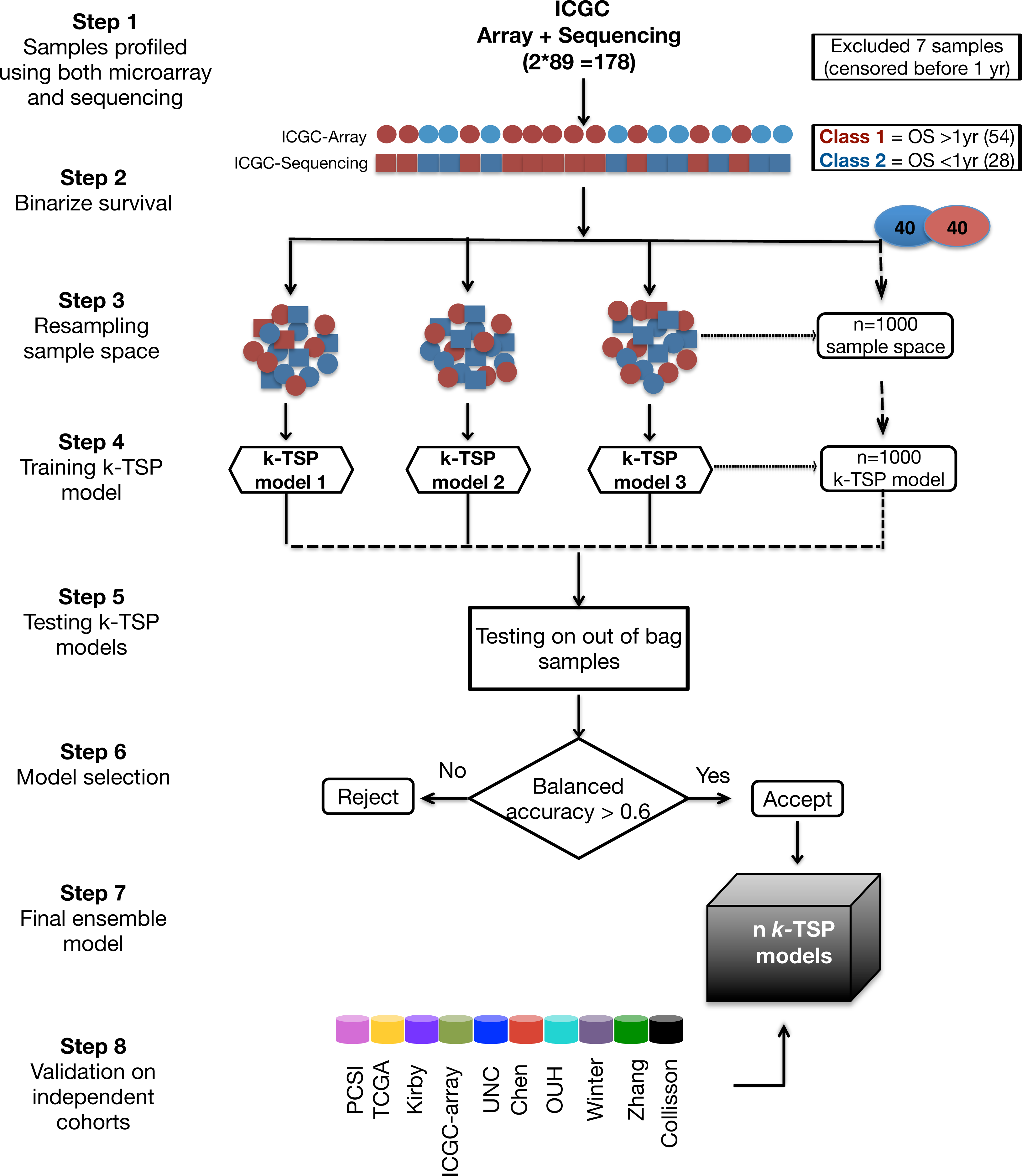
Pipeline showing the approach used for building the Pancreatic Cancer Overall Survival Predictor (PCOSP).

### Datasets

We surveyed the literature and curated 17 datasets including 1,236 PDAC patients from public domain for which transcriptome data of PDAC are available (Supplementary Table S1). We further filtered samples based on the availability of overall survival (OS) and sample size (>10) after dichotomization into high and low survival groups based on an OS cut-off of 1-year (Figure 2). This resulted in a total of four sequencing studies and seven array-based studies providing transcriptomic and clinical data for 1,001 PDAC patients. A total of 12,430 protein-coding genes commonly assessed across all the cohorts were used for further analysis. Clinicopathological features of all the cohorts are presented in Supplementary Table S2. The different cohorts had similar clinical presentation, and were treated with curative surgery followed by adjuvant chemotherapy. Two third of the patients completed multimodal treatment (i.e., surgery and adjuvant chemotherapy; Supplementary Table S2).

**Figure 2:**
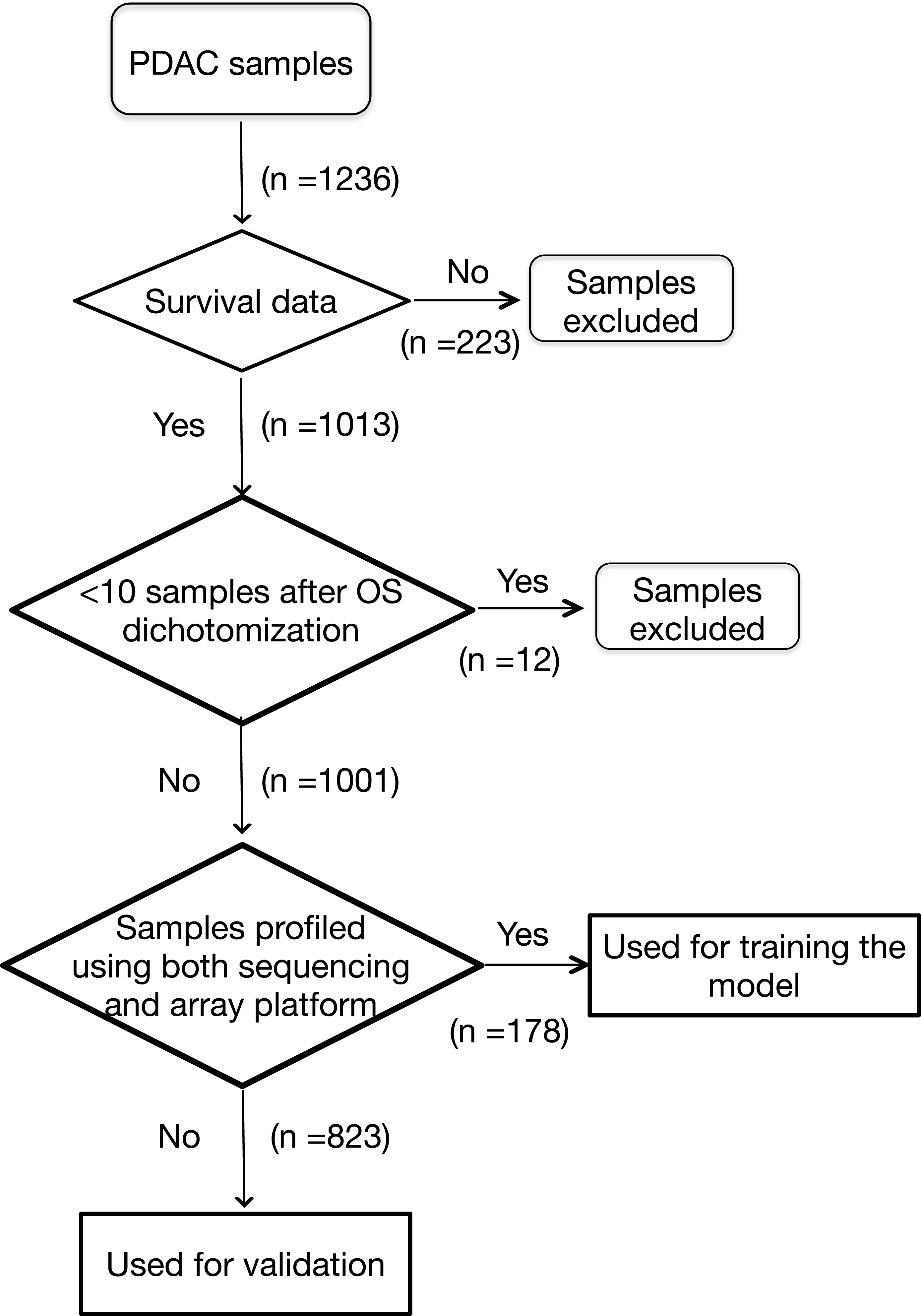
Flowchart showing the inclusion criteria for the pancreatic adenocarcinoma samples.

### Prognostic model

To develop a robust predictor for early death, we used the gene expression profiles of 89 PDAC patient samples whose tumors have been profiled using both microarray and sequencing platforms within the International Cancer Genome Consortium (ICGC) cohort. Approximately half of the patients of the training cohort which where eligible for surgery relapsed within 1 year, we used this threshold to predict PDAC patients with high risk of early death (≤1 yr) post surgery. We excluded 7 samples from the training cohort as these patients were censored before one year of follow-up.

To make gene expression profiles comparable between the training and validation sets, we transformed the original gene expression profiles into binary gene pair barcodes. The advantages of considering pairs of genes with a binary value (“1” if expression of gene *I* > gene *j*, “0” otherwise) are; (*i*) it transforms the feature space in a way that mitigates platform biases and potential batch effects; (*ii*) it makes the model robust to any data processing that preserves the gene order^25,26^. We implemented k-Top Scoring disjoint Pairs (k-TSP) classifier predictor^27^ using the Wilcoxon rank sum method as filtering function in the *SwitchBox* package (version 1.12.0)^28^.

The decision rules are based on the relative ordering of gene expression values within the same sample, where the *k* top scoring gene pairs are used to build the classifier. The samples were resampled 1000 times, where 40 samples from each group were selected in each run to build a *k*-TSP model and the model was further tested on the 49 out-of-bag samples. The models were selected if the balanced accuracy was above 0.6 else the model was rejected. We then froze the parameters of the predictive model and validated it in the remaining compendium of independent datasets. The class probability of the sample was calculated as the frequency of sample predicted as one class divided by the total number of models.

### Random classifiers

To test whether the prognostic value of the PCOSP model could be achieved by random chance alone, we implemented two permutation tests. To test whether the gene expression profiles were associated with survival, we shuffled the actual class labels while maintaining the expression values. To test whether the gene pairs selected in the PCOSP model were robustly associated with survival, we randomly assigned genes to the k-TSP model and assessed its prognostic value. Both procedures were performed 1000 times. As a pre-validation set we compared the balanced accuracy of all the 1000 random models generated using both the approaches to PCOSP using the Wilcoxon rank sum test. Further, we trained the k-TSP classifier models from both approaches in the same way as we built our consensus PCOSP model. We then froze the parameters of the prognostic model and validated it in the compendium of independent datasets, and compared the meta-estimates for both the models against the PCOSP model.

### Early death prediction

The meta analysis was performed for the PDAC sequencing cohorts, PDAC array-based cohorts and the overall combined cohorts to assess and statistically compare the performance of the PCOSP. The patient samples were dichotomized into two groups based on the outcome variable (time from surgery to death ≤ 1 year). Samples censored before 1 year of follow-up were excluded from the analysis of meta-estimate of the area under the receiver operating characteristics curve (AUROC). The AUROC plots the sensitivity vs. 1-Specificity and is used as a criterion to measure the discriminatory ability of the model^29^. The AUROC was computed using *pROC* package (version 1.10.0), and the p-value was estimated using the Mann-Whitney test statistics estimating whether the AUROC curve estimate is significantly different from 0.5 (random classifier). The meta-estimate of AUROC was estimated using the random effect model^30^ implemented in *survcomp* package (version 1.26.0)^31,32^.

### Survival prediction

Prognostic value and statistical significance of survival difference between the predicted classes were assessed using the D-Index, which is a robust estimate of the hazard ratio comparing two equal-sized prognostic groups^33^. In addition, we used the concordance index (C-index) which estimates the probability that, for a random pair of patients, the PCOSP score for the patient with shorter survival is higher than the patient with longer survival^34^. Both the robust hazard ratio (HR) and the C-index were calculated using the *survcomp* package. The meta estimate of HR and C-index were calculated for the PDAC sequencing cohorts, the PDAC array-based cohorts and the combined PDAC sequencing and array-based cohorts using the random effect model^30^ implemented in *survcomp* package.

### Subtyping of PDAC cohorts

The PDAC cohorts were classified into basal and classical transcriptomic subtype using the Moffitt classification method^13^.

### Clinicopathological features based model to predict early death

The clinical model was built by fitting the logistic regression model using common clinicopathological features available from all the cohorts, i.e., age, gender, TNM status and tumor grade.

### Gene set enrichment analysis

To categorize genes in the PCOSP, we performed gene set enrichment analysis using RunGSAhyper function implemented in piano package (version 1.16.4)^35^. The genes selected in the PCOSP model were compared against Gene Ontology (GO) gene sets, canonical pathways and hallmark gene sets using as background the protein-coding genes commonly assessed across the gene expression profiling platforms in our data compendium. Enrichment p-values were corrected for multiple testing using the false discovery rate approach (FDR < 5%)^36^.

### Comparison to existing classifiers

We calculated the Birnbaum signature scores^22^ and Chen signature scores^23^ using the published coefficients of the 25 and 15 classifier genes, respectively, as weight parameter in the *sig.score* function implemented in the *genefu* R package (version 2.10.0)^37^. The Haider signature scores were used as courtesy of the author^24^. The C-index and HR were computed for the three classifiers using eight validation cohorts excluding the cohorts used for training by PCOSP and other classifiers in comparison. Further, we compared the meta-estimates of C-index of each classifier with PCOSP at P<0.05 (one-sided t-test) as implemented in *survcomp* package.

### Research reproducibility

Our code and documentation are open-source and publicly available through the PDACSurv GitHub repository (github.com/bhklab/PDACsurv). A detailed tutorial describing how to run our pipeline and reproduce our analysis results is available in the GitHub repository. A virtual machine reproducing the full software environment is available on Code Ocean. Our study complies with the guidelines outlined in^38–40^. All the data are available in the form of R package MetaGxPancreas (http://bioconductor.org/packages/MetaGxPancreas/).

## Results

### Overall survival predictive model

To predict the patients with early death (≤ 1 year after surgery), the PCOSP model was trained on the 89 ICGC cohort samples profiled using both microarray and sequencing transcriptomic profiles (Supplementary Table S1). To develop a predictor that can be applied to multiple profiling platforms, we transformed the gene expression profiles into binary gene pairs (*x*=1 if expression of gene *i*> gene *j*, *x*=0 otherwise) and used these transcriptomic barcodes in an ensemble of 1000 *k-TSP* predictive models. The PCOSP score is subsequently calculated using the majority voting rule. We tested the prognostic value of PCOSP score in three independent sequencing cohorts, including the Pancreatic Cancer Sequencing Initiative (PCSI)^41^, TCGA-PAAD^15^ and Kirby^42^ cohorts, and seven independent array-based cohorts composed of ICGC-array (excluding the 89 samples used for training)^43^, UNC^13^, OUH^44^, Chen^23^, Zhang^45^, Winter^46^ and Collisson cohorts^12^(Supplementary Table S1). We first tested the predictive value of early death by calculating the AUROC for each dataset separately (Figure 3A). PCOSP was highly significant overall (AUROC=0.70; P<1.9E-18; Figure 3A), although higher in the datasets generated using sequencing platforms compared to microarrays (AUROC 0.72 vs 0.68 for sequencing and array datasets, respectively). PCOSP was significantly predictive of early death in all cohorts (AUROC∈[0.67,0.76]; P<0.05) except the Winter and OUH cohorts (P>0.48) and was almost significant for the Collisson cohort (AUROC=0.69; P=0.051). To determine whether the early death predictive value of the PCOSP model can be achieved by random chance alone, we first computed meta-estimates of AUROC by randomly shuffling the class labels (early deaths) 1000 times and applying the same training procedure used for the PCOSP model. We observed that the gene expression profiles were significantly associated with survival as none of the random models could yield a predictive value greater or equal to PCOSP (p<0.001; Supplementary Figure S1A). We further tested whether the gene pairs selected in the PCOSP model were robustly associated with early death events, by randomly assigning genes to the PCOSP model. Again, we observed that the genes selected in PCOSP yielded significantly more predictive information than the models comprised of random genes (p<0.001; Supplementary Figure S1B), supporting the biological relevance of the PCOSP gene set.

**Figure 3.**
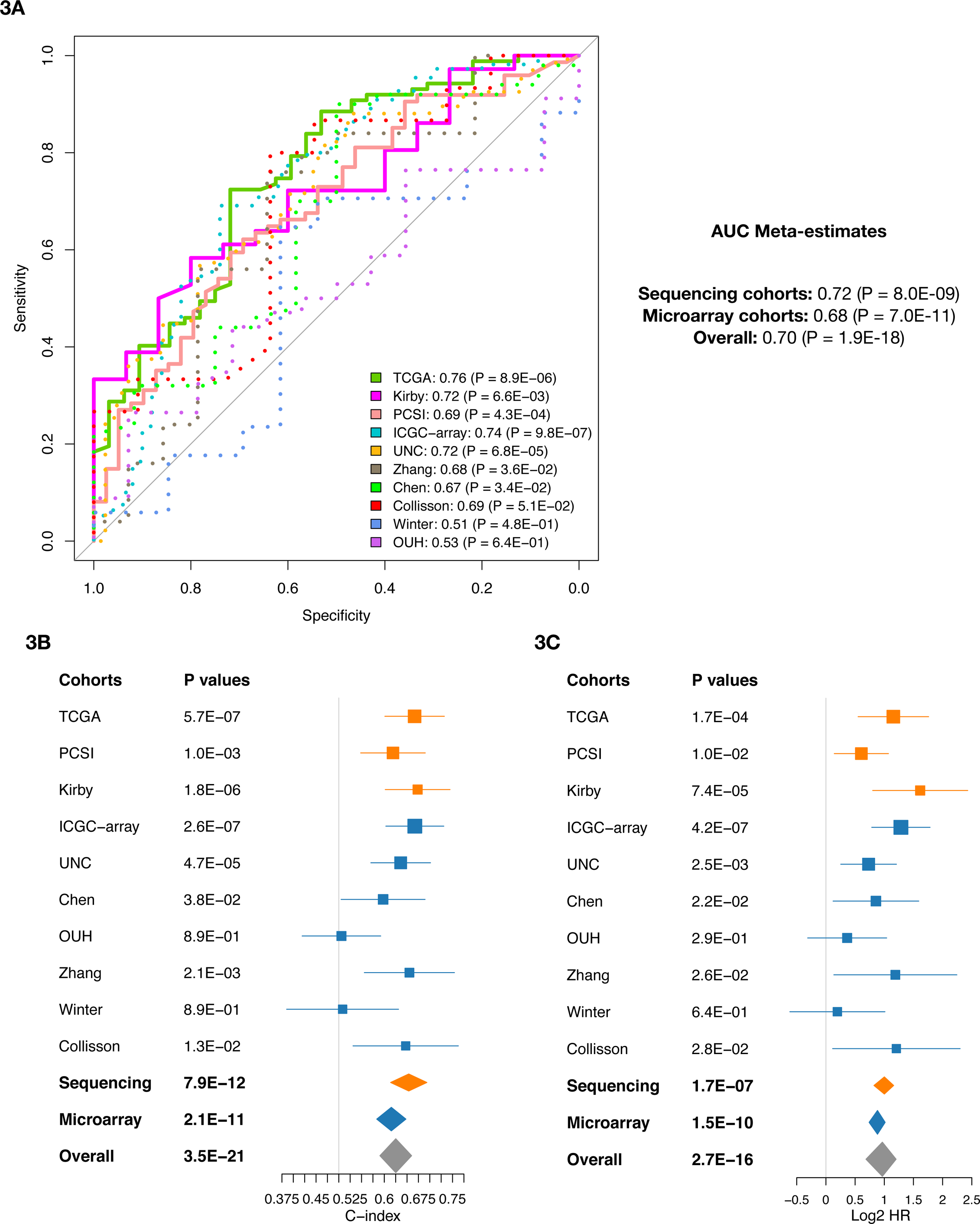
Predictive value of PCOSP for early death and overall survival. (**A**) Area under the ROC curves for all the cohorts and the meta estimates for sequencing cohorts, array-based cohorts and for both the platforms combined. (Forestplot reporting (**B**) the concordance indices (C-index) and (**C**) the hazard ratio (HR) for all the cohorts and the meta estimates for sequencing cohorts (orange), array-based cohorts (blue) and for both the platforms combined (grey).

### Prognostic relevance of the PCOSP model

To assess the prognostic value of the PCOSP model, we calculated the C-indices and HR using the overall survival data for all the cohorts. The C-index is significant overall (C-index=0.63, P=3.5E-21; Figure 3B). In agreement with the results of early death prediction, the PCOSP prognostic value was higher for the sequencing datasets when compared to the arrays (C-index=0.65 vs 0.61 for sequencing and array datasets, respectively; P<2.1E-11; Figure 3B). Similar to the C-index, the PCOSP HR was strong and significant overall (HR =1.95, P=2.7E-16; Figure 3C), and stronger for the sequencing datasets (HR = 2.24 vs 1.83; Figure 3C). To assess whether the prognostic value of PCOSP depends on PDAC molecular subtypes, we stratified PDAC samples into the basal and classical subtypes and calculated meta-estimates of C-index and HR (Supplementary Figures S2A and S2B). We found that PCOSP was prognostic in validation cohorts independently of molecular subtypes. We further tested whether PCOSP prognostic value was complementary to clinicopathological parameters and molecular subtypes by fitting both a multivariate Cox proportional hazard model to predict survival and a logistic regression model to predict binary outcome (death >1yr or not) (Supplementary Table S3).

### Clinicopathological model to predict OS

Patient-specific clinicopathologic features were available for the PCSI, ICGC-sequencing, ICGC-array, TCGA and OUH cohorts. The common variables available were age, gender, TNM status and tumor grade. We fitted the logistic regression model using these clinicopathological features to predict early death of PDAC patients. The clinicopathological model was significant overall with however a weak predictive value (C-index=0.55; P=0.04; Figure 4A). Contrary to PCOSP, the clinicopathological model was not predictive in the sequencing cohort (C-index=0.53 vs 0.58 with P=0.25 vs 0.03 for the sequencing and the array datasets, respectively; Figure 4A). Only nodal status, tumor grade and molecular classes were significant in the univariate analysis (Supplementary Table S3). We compared the prognostic value of the clinicopathological model against PCOSP (Figure 4B,C). PCOSP was significantly more prognostic than the clinicopathological model (one-sided t-test P < 0.01; Figure 4D).

**Figure 4.**
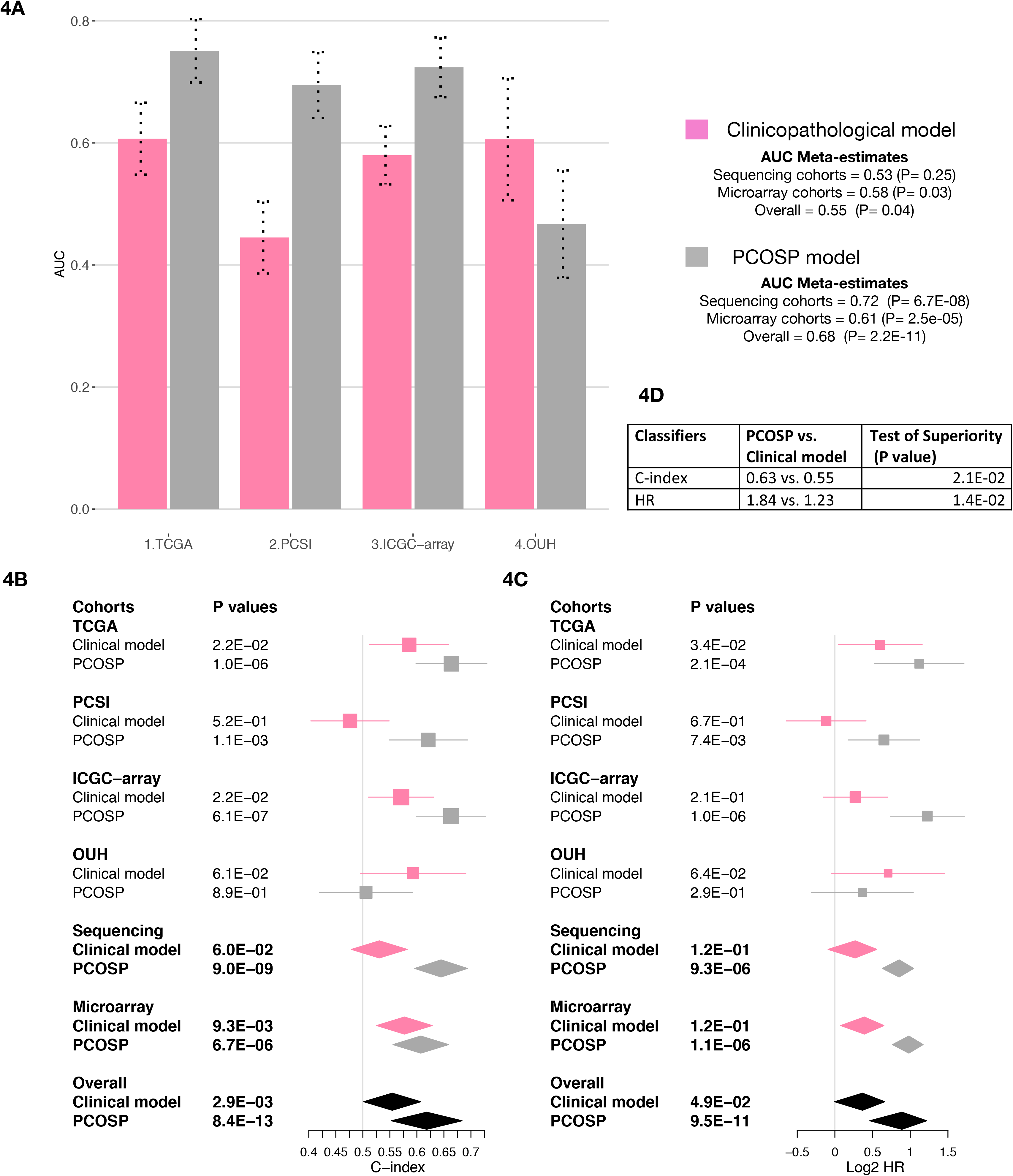
Comparison of the prognostic value of the clinicopathological model and PCOSP. (**A**) Barplot reporting the AUROCs for the clinical model and the PCOSP model. (Forestplot reporting the the (**B**) concordance index (C-index) and (**C**) Hazard ratio (HR) of validation cohorts computed using PCOSP, and clinicopathological model.

### Comparison with published prognostic models

We compared the prognostic value of PCOSP to three published PDAC prognostic models, referred to as Birnbaum^22^, Chen^23^ and Haider^24^. The overall prognostic value of the three published models was significant (P<8.5E-7; Figure 5A,C). PCOSP significantly outperformed published prognostic models in all cases (P<0.05, Figure 5C,D); except for the HR of the Chen classifier where the superiority of the PCOSP prognostic value showed a trend to significance (one sided t-test P=0.10).

**Figure 5.**
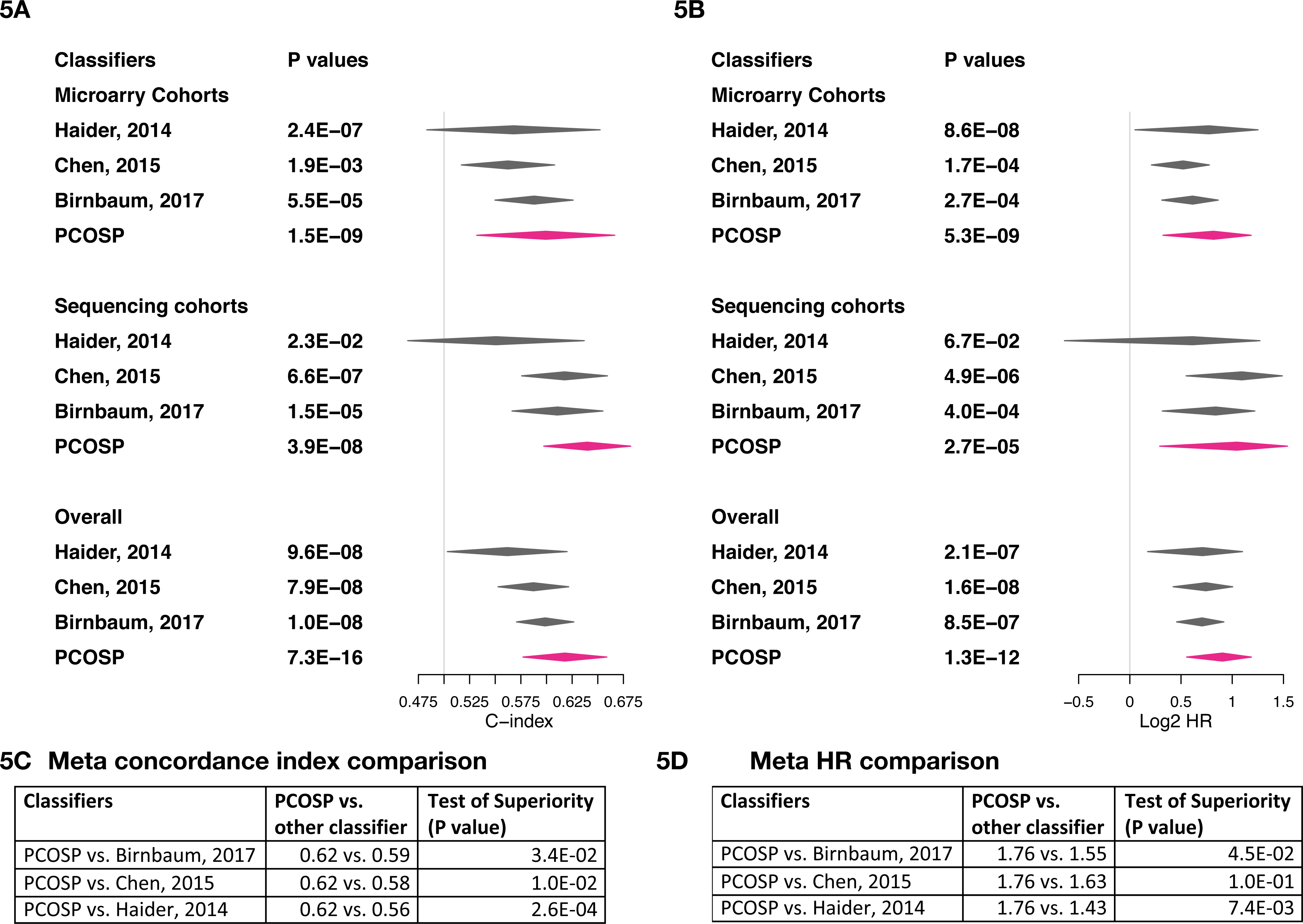
Comparison of existing classifiers with PCOSP. The forestplot reports the meta-estimate of (**A**) concordance indices (C-index) and (**B**) hazard ratio (HR) for PCOSP and existing classifiers.

### Pathway analysis of prognostic genes

Gene enrichment analysis for PCOSP signature genes was performed using hypergeometric test using the hallmarks gene sets, GO molecular function, GO cellular component terms and canonical pathways in MSigDb^47^ with the 1,070 unique genes from the PCOSP model. The Extracellular matrix (ECM), Epithelial Mesenchymal transition (EMT) and hedgehog signalling pathway genes were enriched in the PCOSP model at false discovery rate (FDR) <5%. The complete list of GO terms and pathways significantly enriched in the PCOSP model are listed in Supplementary Table S4A-4D.

## Discussion

We performed a meta-analysis of the transcriptomic profiles of 1,236 PDAC patients and developed PCOSP, a new prognostic model to identify patients with high risk of early death after surgery. The model is built from a unique set of 89 patients profiled using both array-based and sequencing platforms, and validated on a compendium of ten independent datasets, including 823 patients. The prognostic value of the PCOSP model was highly significant for both early death (≤1 year) and overall survival (P<0.001; Figure 3).

Contrary to published prognostic signatures fitted on small number of samples and lacking validation in large independent datasets^17–21^, PCOSP has been trained and validated on a large compendium of datasets. Comparison of PCOSP with existing classifiers^22–24^ showed that the Birnbaum, Chen and Haider models yielded significant (P<0.8.5E-7) but significantly weaker prognostic value than PCOSP (Figure 5C,D). Importantly, PCOSP performs significantly better than existing classifiers for both microarray and sequencing platforms, likely due to simplifying the continuous expression space into binary pair barcodes. This enables PCOSP to be used as a single sample predictor robust to profiling platforms, potential batch effects and normalization methods compared to other classifiers.

Comparison of PCOSP against known prognostic clinicopathological variables showed that PCOSP outperformed the clinicopathological model in predicting early death (Figure 4). PCOSP prognostic value was significant, even after adjusting for molecular subtyping (classical vs basal) and clinicopathological parameters (age, sex, TNM status, differentiation grade of tumor and molecular classes); Supplementary Figure S2A,B and Supplementary Table S3).

The PCOSP model incorporates 2,619 unique gene pairs, totalling 1,070 unique genes. Functional analysis of 1,070 genes showed enrichment of Hedgehog signalling, ECM and EMT pathway. Numerous studies have suggested the involvement of EMT in invasion and metastasis of PDAC^48^. EMT enhances cell motility through loss of cell-cell adhesion, escaping from extracellular matrix and overcoming the apoptosis process^48^. The ECM and EMT pathways are not only associated with the metastatic spread of tumor but also with chemoresistance that leads to worse survival^49^.

PDAC is a heterogeneous and genetically highly complex disease, supporting the molecular^13,14^ and morphological^50^ characterization of a given tumor as an important cornerstone for the development of future therapies. We provide the largest compendium of 17 PDAC datasets as a gold standard for future PDAC analyses. The new meta-analysis framework implemented in PCOSP maximizes robustness and performance across the cohorts. In order to implement PCOSP as a clinical assay, we tested different feature set sizes for the k-TSP models and compared the performance of the reduced models. We achieved accuracy comparable to the 1,070 gene-PCOSP model by including only 256 unique genes, supporting the potential of a smaller PCOSP based useful in the clinic (Supplementary Figure S3). Endoscopic ultrasound (EUS) biopsies could be utilized prior to curative surgery to estimate the prognosis of PDAC patients using PCOSP. This may assist clinicians in making postoperative treatment decision, i.e., palliative therapy, standard adjuvant chemotherapy or alternative chemotherapy regimens, which holds the potential to improve the overall survival of the patients.

The current study has potential limitations. First, there are inherent tumor sample collection biases as the different datasets were collected and sampled at different centers. The levels of tumor cellularity varied highly across cohorts as PCSI and Collisson datasets were generated using laser microdissection prior to sequencing, Kirby and Chen datasets were macrodissected, while TCGA, ICGC, OUH, Zhang and Winter datasets used bulk tumors for profiling. Second, the transcriptomic profiles in our data compendium were generated using different gene expression profiling technologies for sequencing (Illumina HiSeq 2000/2500) and microarray platforms (Agilent, Affymetrix, and Illumina). Third, all samples were normalized using the published processing methods, which depend on the profiling platforms (Supplementary Table S2). Despite these limitations, PCOSP yielded robust prognostic value across the heterogeneous datasets, indicating that the gene expression barcode transformation is robust to the inevitable biases present in large meta-analyses.

The lack of available clinical and treatment information across the cohorts is also a limiting factor in our meta-analysis. However, comparison of cohort specific clinical information for the cohort were not significantly different across the cohorts (Supplementary table S2). During the time period of sample collection, standard of care treatment for PDAC was curative-intent surgery followed by adjuvant chemotherapy with gemcitabine or 5-FU. New approaches using doublet and triplet chemotherapy regimens are now standard of care in the palliative setting and randomised trials using these agents in the adjuvant setting will be reported shortly. Neoadjuvant therapy is also being evaluated in many centres. Thus, heterogeneity in treatment is expected within and between different cohorts, we will need to test our PCOSP model using new clinical datasets, or preferably within the context of randomized trials.

## Conclusion

We leveraged the largest compendium of PDAC transcriptomes to develop PCOSP, a prognostic model identifying PDAC patients at high risk of early death independently of, and superior to, clinicopathological features and molecular subtypes. PCOSP may be useful in the clinical setting as a single sample classifier to identify patients who could be at higher risk of early death following surgery and adjuvant chemotherapy, potentially facilitating treatment decisions, including the use of neoadjuvant chemotherapy or alternate treatment strategies.

## Acknowledgements

This study was conducted with the support of the Ontario Institute for Cancer Research (OICR, PanCuRx Translational Research Initiative) through funding provided by the Government of Ontario (Ministry of Research, Innovation, and Science), and a charitable donation from the Canadian Friends of the Hebrew University (Alex U. Soyka). V.S was supported by grants from The Radium Hospital Foundation, Oslo University Hospital, and the PanCuRx Translational Research Initiative at the OICR. B.H.K was supported by the Gattuso Slaight Personalized Cancer Medicine Fund at Princess Margaret Cancer Centre, the Canadian Institutes of Health Research, the Natural Sciences and Engineering Research Council of Canada, and the Ministry of Economic Development and Innovation/Ministry of Research & Innovation of Ontario (Canada). We thank all the patients who participated in the study.

## SUPPLEMENTARY FIGURES

**Supplementary Figure S1:**
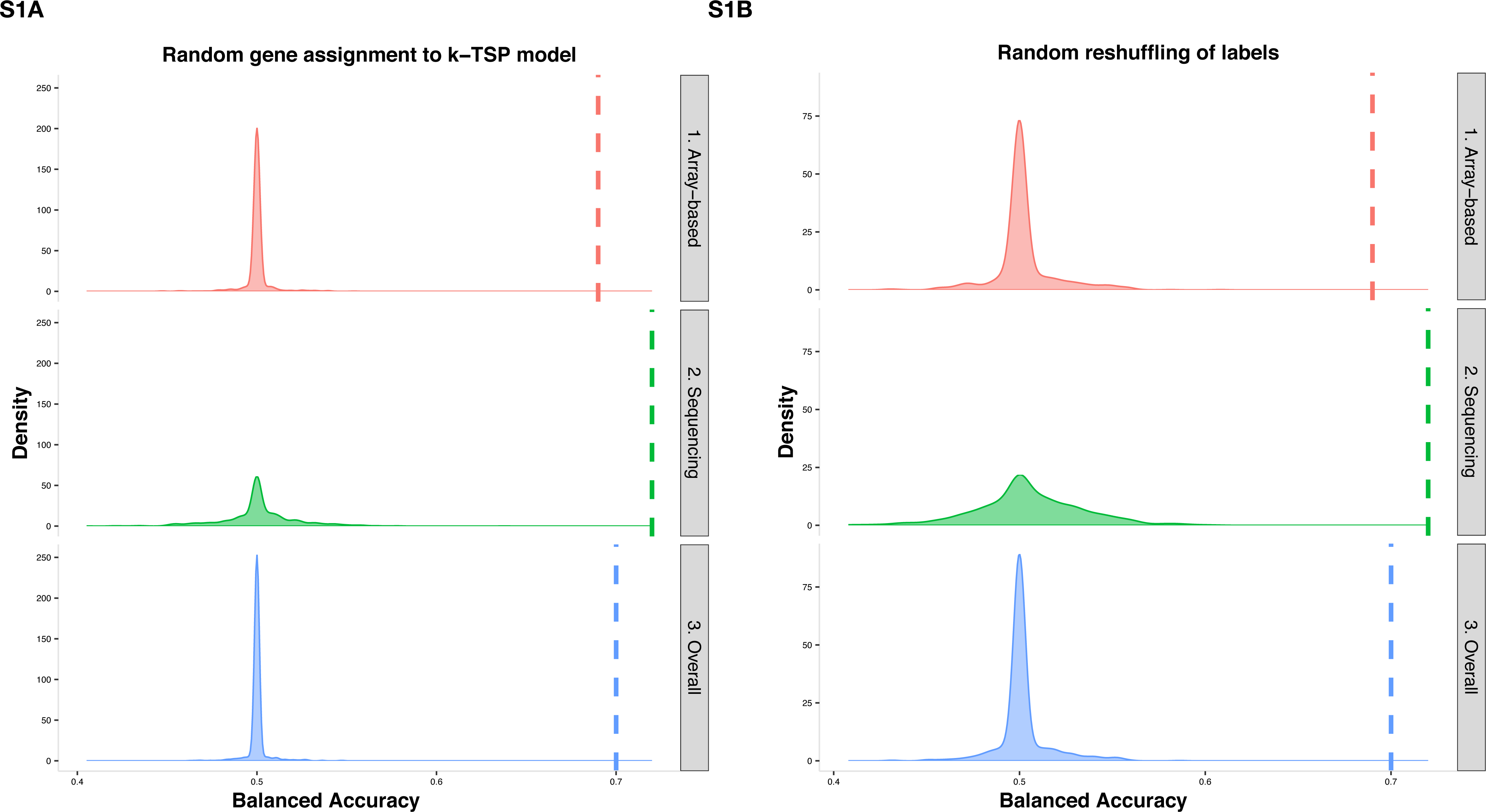
Density plot showing the distribution of balanced accuracy for random models. Distribution of meta-estimates of 1000 models generated using (**A**) random reshuffling of labels and (**B**) random assignment of genes to TSP models. The meta-estimates were independently calculated for all the cohorts combined, sequencing cohorts and array-based cohort. The pink, green, blue dashed lines represent meta-estimate of AUROC from PCOSP model for overall, sequencing and array-based cohorts respectively.

**Supplementary Figure S2:**
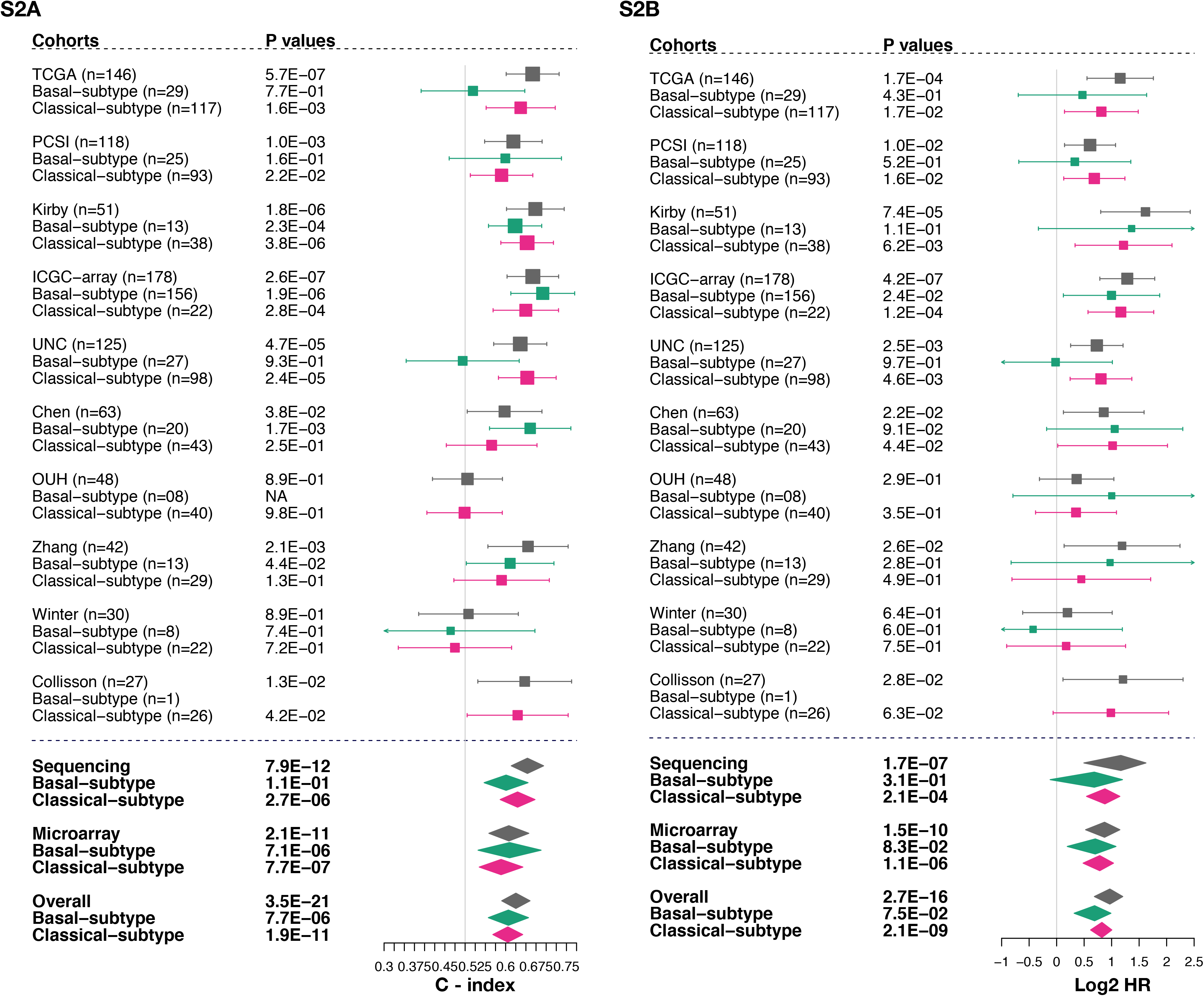
Forestplot of (**A**) concordance index (C-index) and (**B**) hazard ratio (HR) for all the cohorts divided based on the molecular subtypes. The grey, green and pink color in the forestplot depicts meta-estimate of C-index for overall cohort, the basal subtype and the classical subtype of the cohorts, respectively.

**Supplementary Figure S3:**
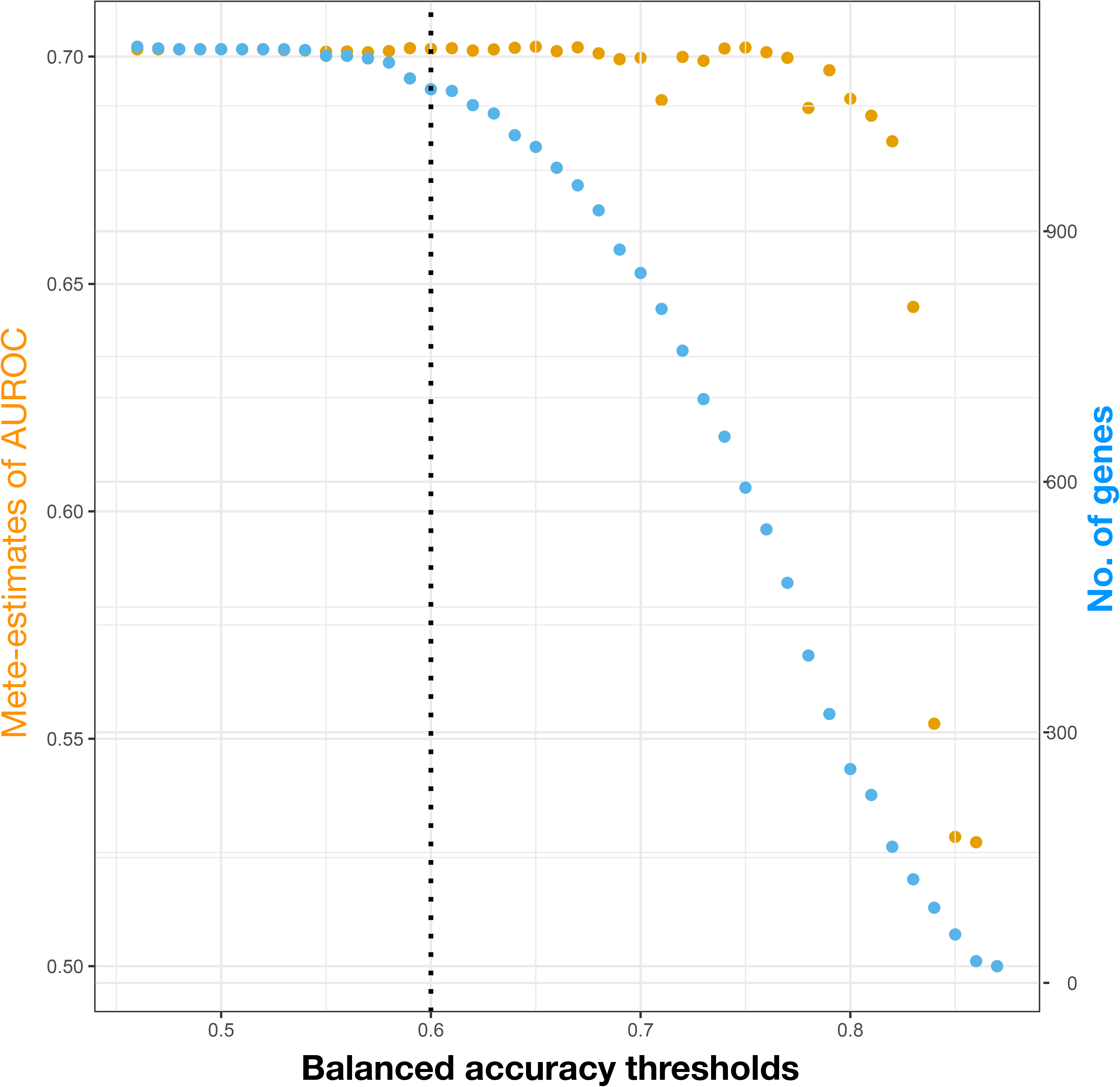
The scatterplot shows the meta-estimate of AUROC (orange) and total number of unique genes (blue) in the PCOSP model at different balanced accuracy thresholds. The threshold used in the PCOSP is marked as dashed line at 0.6.

## SUPPLEMENTARY TABLES

**Supplementary Table S1:** The table shows the datasets used in the project for meta-analysis.

**Supplementary Table S2:** The table shows the clinicopathological information of the validation cohorts used in the analysis.

**Supplementary Table S3:** Univariate and multivariate regression analysis.from (**A**) logistic regression model to predict early death (death >1 yr or not), and (**B**). the Cox regression model using clinicopathological features, molecular subtypes and PCOSP model probabilities for validation cohorts.

**Supplementary table S4:** The table shows the pathways overrepresented in the PCOSP model genes using (**A**) hallmark gene sets, (**B**) canonical pathways, (**C**) GO-molecular function term (and (**D**) GO cellular component terms from MSigDB.

